# Mapping lineage-resolved scRNA-seq data with spatial transcriptomics using TemSOMap

**DOI:** 10.1101/2024.10.31.621331

**Authors:** Xinhai Pan, Alejandro Danies-Lopez, Xiuwei Zhang

## Abstract

Spatial transcriptomics (ST) has revolutionized the study of cell spatial organization and cell-cell interactions. However, current ST technologies face limitations such as lower gene coverage and spatial resolution compared to single-cell RNA sequencing (scRNA-seq). Integrating scRNA-seq with ST can address these issues by mapping single cells onto spatial data, thereby inferring their spatial coordinates. During tissue formation, cells derived from the same ancestor often remain spatially proximate, making lineage data valuable for cell location inference. Certain single-cell multi-omics technologies including lineage tracing provide paired gene expression and induced or sometic mutation information in single cells, which can be used to reconstruct the cell lineage tree, representing the clonal relationships of cells. To incorporate this information, we developed TemSOMap (**Tem**poral dynamics guided **S**patial **O**mics **Map**ping), which infers the spatial coordinates of cells by mapping a paired gene expression and mutation barcode dataset onto a spatial transcriptomics dataset. TemSOMap infers a cell-to-spot mapping matrix by minimizing a loss function incorporating gene expression, cell lineage and cell location information. We show that TemSOMap more accurately infers the spatial location of single cells compared to state-of-the-art baseline methods under various scenarios, using both simulated and real datasets. The resulting lineage-resolved ST data can help us better understand the spatio-temporal dynamics of cells in a tissue. TemSOMap is publicly available at https://github.com/ZhangLabGT/TemSOMap.

## 1 Introduction

Spatial Transcriptomics (ST) has become one of the most widely adpoted technologies in single-cell multi-omics. This technology has enabled the study of the spatial distribution of gene expression patterns within complex biological structures, providing valuable insights into the organization and function of cells in their native environment [2, 8, 32]. The state-of-the-art ST technologies can be summarized into two categories: sequencing-based and imaging-based. The 10x Visium technology [1] is a widely adopted sequencing-based method with reasonable cost. This technology can sequence the whole transcriptome, but the measured gene expression is not at the single-cell level; instead, it measures gene expression profiles at each *spot*, which often covers multiple cells. Stereo-seq [20], on the other hand, can measure sub-cellular resolution spots but still does not provide cell-level gene expression data. For imaging-based methods (such as CosMx [11], 10x Xenium [14, 18], MERFISH [7], STARmap [31]), since gene expression is captured *in situ*, the cell location information can be obtained, but these technologies can only capture up to hundreds of genes at the same time.

To overcome the limitations of the ST technologies, researchers have explored the possibilities of integrating or mapping ST data and other single-cell omics data, and in particular, scRNA-seq data. Methods have been proposed to integrate scRNA-seq and ST data, which serve various objectives including predicting locations of cells in the scRNA-seq data, imputing missing gene expression in the ST data, and estimating cell type proportions of the spots in low-resolution ST data [21, 19]. In this paper, we focus on the task of predicting cell locations from the scRNA-seq data. Although our method can work with ST data from various technologies, we focus on the “low-resolution” spot-based ST data (like those from 10x Visium) when describing our methods, as these data are the most abundant while also being the most challenging to use. Tangram [3] infers a cell-by-spot mapping matrix utilizing gene expression dissimilarity loss between mapped and the reference ST data. SpaOTsc [4] is a method developed based on optimal transport (OT) that infers spatial coordinates for scRNA-seq data. CeLEry [33] uses a variational autoencoder to learn the mapping between scRNA-seq and ST data. Though these methods vary in their methodologies, they share the fundamental ideas of using the expression levels of common genes in both datasets as a bridge to map cells in scRNA-seq data to locations in ST data. However, due to the batch effects between ST and scRNA-seq data, the large search space of potential spatial locations for cells in the scRNA-seq data, and the reduced number of shared features between ST and scRNA-seq data, inferring the spatial locations of cells based solely on gene expression data is still a very challenging problem, and the accuracy of state-of-the-art methods is unknown due to the lack of ground truth data.

We consider that the temporal dynamics of cells are closely coupled with spatial information of cells, therefore, the temporal information of cells can inform spatial locations of cells. In particular, tissues are formed through generations of cell divisions, which is a temporal process. The fundamental concept of this work is that the information on the temporal process can inform the spatial organization of cells in the tissue. For example, cells divided from the same ancestral cells tend to be located closely in space unless the cells migrate to distant locations after division. While the *cell lineage* or *cell clonal* information can be obtained from multiple types of data that contain genetic somatic mutations or induced mutations [17, 30], in this paper, we take advantage of the CRISPR/Cas9-based *lineage tracing* datasets [29, 5, 26], which consist of paired gene expressions and lineage barcodes in single cells. That is, a lineage tracing dataset consists of a scRNA-seq count matrix, and a set of lineage barcodes, each for a single cell. For any two cells, the difference between their lineage barcodes should reflect their distance on the cell division tree. Therefore, the lineage barcodes can be used to reconstruct the cell lineage tree, representing the clonal relationships between cells.

To incorporate the lineage and clonal information of cells to improve the prediction of cell spatial locations, we developed TemSOMap (**Tem**poral dynamics guided **S**patial **O**mics **Map**ping). In TemSOMap, while closer cells on the lineage tree tend to have closer spatial coordinates by the design of the objective function, we are aware that this pattern can be weakened when there are frequent cell migration events, and we have tested the methods under a wide range of cell migration rates. It is worth noting that even when the migration rate is low, it is challenging to obtain accurate cell lineage and clonal information, due to multiple factors including the noise and missing data in the lineage barcode data [28], and the computational intractability of lineage tree inference from lineage barcodes.

Despite the above-mentioned challenges, we show that TemSOMap outperforms other methods in inferring the spatial coordinates of cells, while accurately inferring the spatial distribution of gene expressions, using both simulated and real datasets. TemSOMap is the first method that integrates lineage information with ST data. In addition to improved accuracy of spatial location prediction, applying TemSOMap to real data can output a spatiotemporal map of single cells, which provides both lineage, spatial, and gene expression information of every single cell, and allows for the analysis of the spatiotemporal dynamics of cells. For example, following the lineage from a progenitor, the spatial migration pattern for its descendant cells can be analyzed. Potentially, one can predict the spatial distribution of cells at earlier development time using the spatial coordinates of leaf cells and the lineage tree.

## 2 Methods

### 2.1 Related background

#### 2.1.1 Cell lineage, cell clone, and spatial coordinates

A *cell lineage* refers to the tree representing the cell division history from a root to present-day cells. A *cell clone* refers to a group of cells that are derived from the same ancestor in the cell lineage tree, thus a cell clone corresponds to leaf cells of a subtree in a cell division tree. The spatial coordinates of cells represent the cells’ relative locations in tissue. In a given tissue sample, if cells from the same clone mostly locate closely in the tissue, we say that this tissue has *clonal pattern*.

#### 2.1.2 Biological processes of cells’ spatio-temporal dynamics

The formation and growth of tissue and organs is a crucial question for developmental biology. We consider two key biological models contributing to cell spatial coordinates in a tissue: *cell division* and *cell migration*.

Cell divisions refer to duplicating a parent cell into two daughter cells. Cell migration refers to the process of cells’ movement in tissue, guided by complex biological factors including cell state, cell microenvironment, and overall tissue environment. Cell migration activities can change the clonal pattern in the tissue. Therefore, when testing TemSOMap, we use a variety of migration rates to test the robustness of TemSOMap against cell migrations.

### 2.2 Overview of TemSOMap

As shown in Fig. 1, TemSOMap takes in a lineage tracing dataset with both scRNA-seq count matrix (denoted by *X*) and lineage barcodes, and a ST dataset with both a spot-level gene expression matrix (denoted by *Y*) and spatial coordinates of the spots (denoted by *S*). The lineage barcodes can be used to obtain cell clones(denoted by *c*), where each cell has a clone label, and pairwise barcode distance between cells (denoted by *Z*). The objective is to infer *M*, a cell-by-spot mapping matrix that represents the probability for each cell in *X* being mapped to each spot in *Y*. This definition of the mapping matrix is used in previous work [3]. Then *M*^*T*^ *X* becomes a ST data matrix converted from the scRNA-seq count matrix *X*. Predicted cell coordinates of cells in *X* can be obtained from *MS*. We design an objective function to infer *M*.

**Fig. 1.**
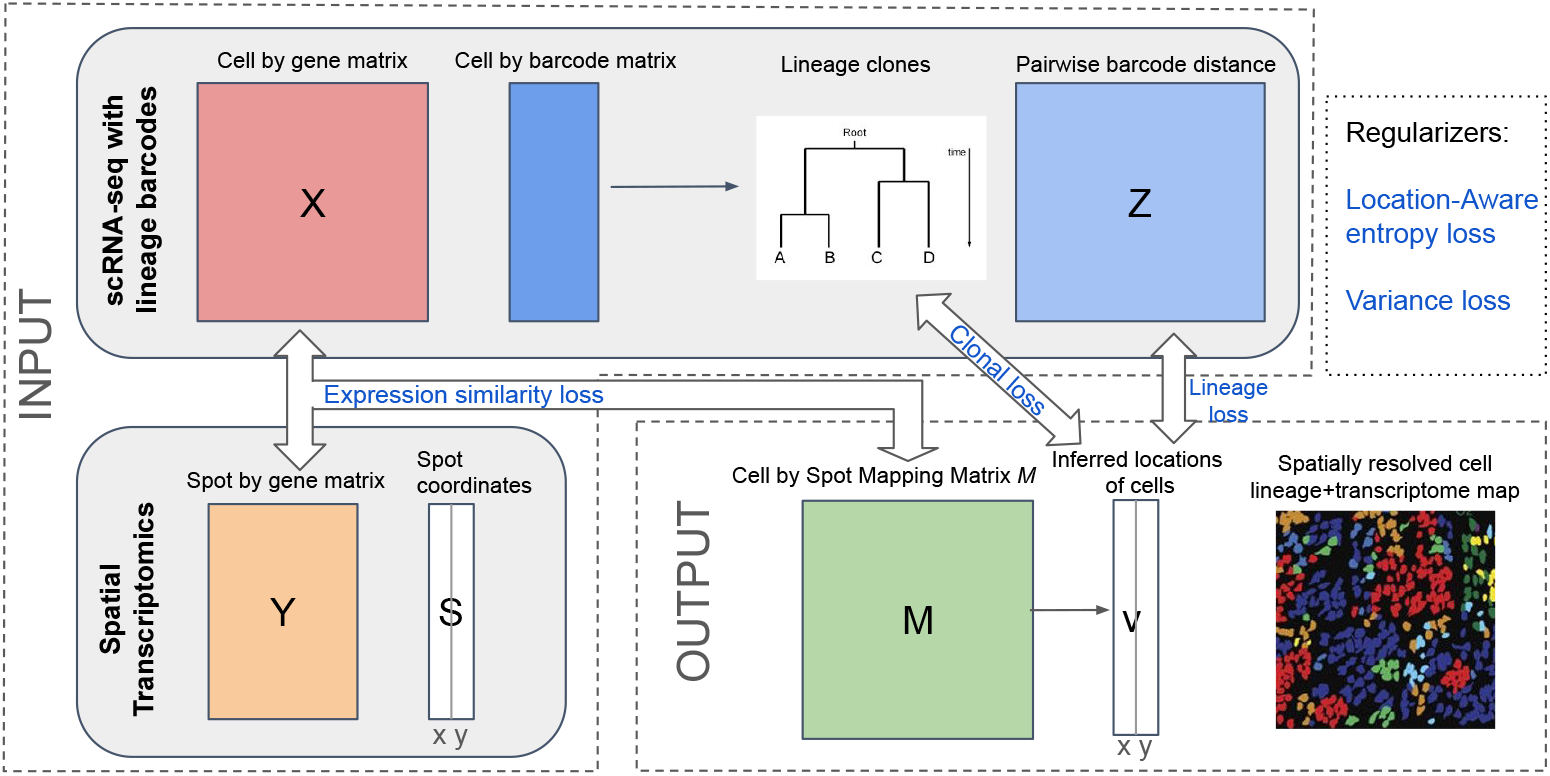
Overview of TemSOMap. The input to TemSOMap includes a scRNA-seq data matrix with single-cell lineage barcodes and a spatial transcriptomic (ST) dataset. TemSOMap outputs a mapping matrix *M*, which can be used to obtain the inferred location of cells. Blue fonts highlight the loss terms (See Methods).

The objective function of TemSOMap consists of three major loss terms and two regularization terms (Fig. 1). The three major terms are: expression similarity loss, which calculates the dissimilarity between *M*^*T*^ *X* (the ST data converted from scRNA-seq) and the given ST data *Y* ; lineage loss, which aims to maintain the similarity of the pairwise distances between cells on the inferred spatial coordinates and on the lineage tree; and clonal loss, which enforces the spatial clustering of cell clones, *i.e*., cells in the same clone should locate close to each other.

We have added regularization terms each aiming at maintaining a desired property of *M*. These are: location-aware entropy loss, which forces the probability distribution of every cell to be sparse and spatially concentrated; and variance loss, which penalizes large variance for the spatial distribution of a cell’s possible coordinates. Then, the total loss is calculated as the weighted sum of the losses described above. Total loss is minimized via stochastic gradient descent to find the optimal mapping matrix *M*. The predicted locations of cells in scRNA-seq data are then obtained from *M*. These steps are described in detail in the following sections.

### 2.3 Notation definitions

The input data to TemSOMap are denoted as follows: 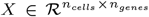 is a cell-by-gene matrix (scRNA-seq count matrix); 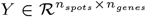, a spot-by-gene matrix representing the ST data; and 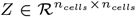, a cell-cell pairwise distance matrix, indicating the distance between single cells in the cell lineage tree. 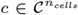 is a vector representing discrete clonal classes for every cell, where 𝒞 = *{c*_1_, *c*_2_, …, *c*_*k*_*}* represent the finite set of possible clones. The spatial coordinates of the spots is denoted as 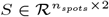 (a spot-by-coordinate matrix).

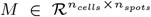 is a cell-by-spot mapping matrix that encodes the soft assignment for each cell to each spot, where 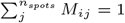 and *M*_*ij*_ *>*= 0 for every cell *i* and spot *j*.

### 2.4 Calculating cell clones and cell lineages from lineage barcode data

Given the barcodes for all cells, to obtain *Z*, we calculate the pairwise weighted Hamming distances between all pairs of barcodes, following practice in existing work [24, 10]. These distance measures between barcodes can capture the geodesic distances between cells lineage and clones. In *Z, Z*_*ij*_ represents the weighted hamming distance of the lineage barcodes between cell *i* and cell *j*. To obtain clone labels *c* for the cells, we use the Neighbor-Joining method [27] to infer a lineage tree *T*, which is a binary tree graph where the leaf nodes are the cells present in the lineage barcode data. Here, the choice of the lineage inference method can be changed and can alternatively include LinRace [24] or Cassiopeia [12]. In TemSOMap, the clonal IDs are learned based on the inferred lineage tree. To obtain balanced clones from the inferred lineage tree, we first re-balanced the lineage tree by rooting the tree at an internal node that has the closest number of cells under the left and right subtrees. Then, the clonal IDs can be obtained by cutting the tree at the specific generation log_2_ *n*_*clone*_ on the re-balanced lineage tree, where *n*_*clone*_ is the number of clones set by users.

### 2.5 TemSOMap mapping algorithm

To constrain values in matrix *M* to be probabilities, we utilize the softmax function to transform a given matrix *M*:

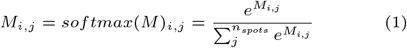

The total loss function of the TemSOMap mapper consists of the following components:

#### 2.5.1 Expression similarity loss

Following Biancalani *et al* [3], we utilize the cosine similarity function to enforce similarity between *M*^*T*^ *X* and *Y*. We apply the cosine similarity function to both rows of *Y*, which correspond to the spot-wise expression patterns (Eq. 2), and columns of *Y*, which correspond to gene-wise expression patterns (Eq. 3). The loss terms can be formulated as follows:

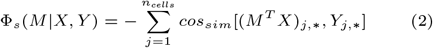

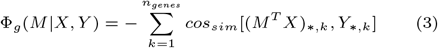

where 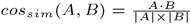.

### 2.5.2 Lineage loss

Using the mapping matrix *M*, we can calculate the pairwise distance matrix between the inferred spatial location of cells. The inferred spatial location of cells can be calculated as 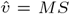, where for each cell, its inferred location is the weighted mean of all locations, with weights being probabilities in *M*. Therefore, the pairwise Euclidean distances between inferred locations of cells can be represented as follows:

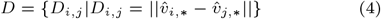

and the lineage loss is the MSE loss between the inferred pairwise distances of cells and the pairwise Hamming distances of cells’ lineage barcodes (with normalization):

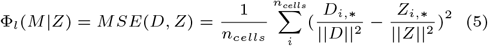

Although the Euclidean distance between cells in space does not perfectly reflect the geodesic distance of cells on the cell lineage tree, it outperformed the inferred geodesic distance from the spatial coordinates in our tests.

#### 2.5.3 Clonal loss

The clonal loss considers cell clones, where cells are assigned clonal labels (described in Sec 2.4) that are similar to cluster labels. In contrast to lineage loss, where all pairwise distances between cells are considered, in the clonal loss, there are two types of distances: intra-clone distance and inter-clone distance. A vector 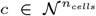 represents the clonal ID for the cells. Considering that cells in the same clone should be located close to each other in space, the clonal loss can be calculated as follows:

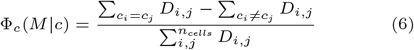

where we try to minimize intra-clone distances and maximize inter-clone distances. The clonal loss is complementary to the lineage loss, and can be more robust to errors in the inferred cell lineage tree.

#### 2.5.4 Location-aware (LA) entropy loss

While the mapping matrix *M* indicates soft assignment, we enforce an entropy loss on the inferred probability distribution of each cell to promote sparsity. Furthermore, we want the mapping matrix to better reflect realistic cell-spot mapping relationships, that is, a cell’s probability among spots should be spatially concentrated. Therefore, we first smooth the mapping matrix using a Gaussian kernel based on the pairwise distances between spots,

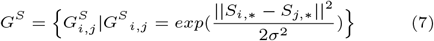

and then calculate the entropy on the smoothed mapping matrix *M*_*s*_, which can be formulated as follows:

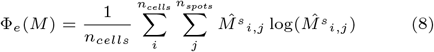

where 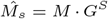, indicating the smoothed mapping matrix.

#### 2.5.5 Variance loss

Each row of the *M* matrix represents a probability distribution among spots for each cell. For each cell, it is desired for spots with high probabilities of including it to be located closely in 2-D space. This can be achieved by minimizing the following weighted location variance that we designed. Given the mapping matrix *M* and the spot coordinates *S* = (**x, y**), indicating the two axes for the coordinates, we calculate the variance loss as the average weighted variance of the spatial distribution of all cells:

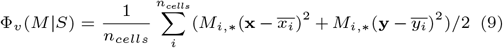

where 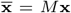 and 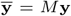, representing the weighted mean locations of cells on *x* and *y* axes, respectively.

#### 2.5.6 Total loss, hyperparameters and optimization

The total loss is therefore calculated as:

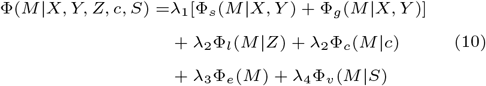

where each loss is weighted using a hyperparameter (*λ*_1−4_). We use a default set of hyperparameters without tuning for the tests we run in this work. The default values are chosen so that the loss terms are of similar magnitude while prioritizing fitting the gene similarities and lineage losses. Details on the list of hyperparameters, their default settings, and rationals for choosing the default setting are in Supp. Info. Sec. 3.2. The minimization of the total loss is achieved via the Adam optimizer using the Pytorch library.

Once the final *M* is calculated, the inferred coordinates of a cell *i* is calculated as

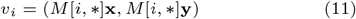

which is the weighted mean of all locations, with probabilities in *M* as weights.

### 2.6 Simulating lineage-resolved single-cell spatial transcriptomics

To benchmark TemSOMap’s performance and compare with other state-of-the-art methods, ground-truth information is needed about cells’ lineage identity, spatial coordinates, and gene expression. Due to the lack of real datasets that contain ground truth information for all three modalities, we developed SpaTedSim, a computational simulation framework that generates cells’ gene expressions, spatial coordinates, and lineage barcodes simultaneously. The simulation process is based on a ground-truth cell division tree, and along the generations of cell divisions, we simulate the lineage barcodes and gene expressions based on our previously published method, TedSim [25] (Supp. Fig. 1a). TedSim can simulate realistic cell state changes on the cell division tree to account for the inconsistency between lineage proximities and expression similarity. Moreover, TedSim generates realistic lineage barcodes with barcode saturation, redundancy, and dropouts. In SpaTedSim, we simulate cells’ movement in space from cell division and cell migration. For cell division, starting with the root cell, SpaTedSim iteratively generates the daughter cells’ (cell *u* and cell *v*) initial coordinates given the parent’s coordinates (*x*_*p*_, *y*_*p*_):

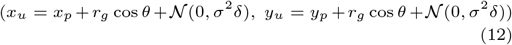

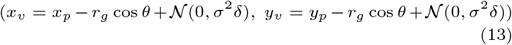

where *r*_*g*_ is the division radius, which depends on the current generation of divisions. The more the cells divide, the smaller the division radius becomes, indicating less freedom of movement when the tissue becomes more crowded and differentiated. *θ* determines the angle of the dividing cells, which leads to opposite moving directions for the two daughter cells. Lastly, 𝒩 (0, *σ*^2^*δ*) represents the 2-D Brownian motion term (Gaussian random walk), where *δ* is the step size and *σ* is the standard deviation.

After cell division, a cell’s spatial coordinates can further change due to cell migration. In real tissues, cell migration is a complex biological process, controlled by various intracellular and extracellular factors. In SpaTedSim, the migration behavior of cells is dependent on their cell types. Given the spatial coordinates of cells of the same cell type, *v* = (*v*_1_, *v*_2_, …, *v*_*n*_) ∈ *R*^*n×*2^, we first estimate the spatial density of the cell type:

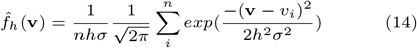

To simplify, we set *σ* = 1, and *h* is the bandwidth hyperparameter defined in density estimation. After each generation of cell divisions, the Gaussian density estimation is performed on every cell type, and cell migration is the process of migrating a cell’s coordinates to a nearby location based on the cell type density map. For a cell’s coordinates after cell division, *v*, and a migration radius *r*_*m*_, the migrated coordinates, *v*^*′*^, are sampled from the cell type densities:

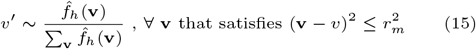

Lastly, after generating all spatial coordinates of the leaf cells, we perform an additional postprocessing step to make the overall coordinates more realistic; that is, cells should have a relatively uniform spatial distribution in the simulated tissue region. It is worth noting that this postprocessing step can weaken the clonal pattern in the data, which leads to increased difficulty for TemSOMap. More details about the simulation algorithms and parameter settings are in Supp. Info. Sec. 2.

## 3 Results

### 3.1 Evaluating TemSOMap on synthetic datasets using SpaTedSim

To quantify the performances of TemSOMap and baseline methods, we use datasets simulated by SpaTedSim. SpaTedSim generates scRNA-seq data and ST data with ground truth cell locations in the scRNA-seq data, and it can vary the amount of batch effects between scRNA-seq and ST data (Supp. Fig. 1b, Supp. Info. Sec. 2.2). By varying migration rate, SpaTedSim can generate datasets with different spatial clustering patterns in terms of cell types or clonal IDs (Supp. Fig. 1c). With a lower migration rate, the cell migration occurs less, and the cells will show better clonal clustering pattern spatially. On the contrary, a higher migration rate results in a better cell-type clustering pattern. To test the performances of TemSOMap, we mask the spatial coordinates of single cells and reconstruct them using spot-level spatial transcriptomics and paired gene expressions and barcodes. An example of visualization of TemSOMap’s reconstructed spatial map of single cells is shown in Fig. 2a, in comparison with the ground truth.

**Fig. 2.**
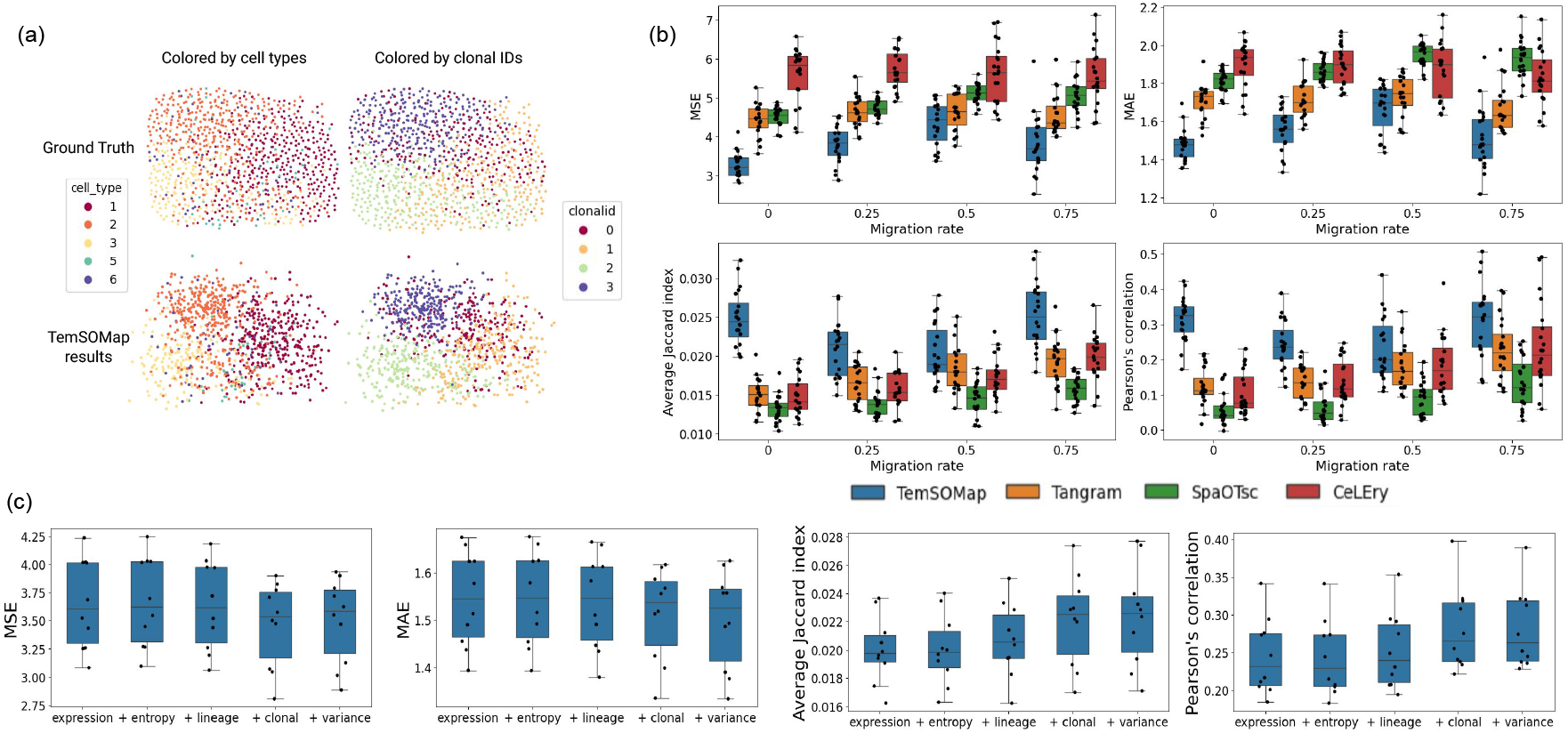
TemSOMap results on SpaTedSim-simulated data. (a) Visualization of the spatial coordinates of single cells, comparing ground truth and TemSOMap inferred results. Left column: cells colored by cell types; right column: cells colored by clonal IDs. The migration rate is 1 for this plot. (b) Comparisons of TemSOMap and baseline methods on SpaTedSim-simulated datasets (1,024 cells), with varying migration rates (0-0.75). For MSE and MAE, lower values mean better accuracy. For the Average Jaccard Index and Pearson’s Correlation, higher values mean better accuracy. No dropouts are added to the lineage barcode data in these results and each migration rate has 20 datasets. (c) Ablation test results of TemSOMap on SpaTedSim datasets with migration rate 0.25. Each box adds one loss term in TemSOMap and includes 10 datasets.

We compare TemSOMap with existing methods that estimate the spatial location of cells: Tangram, CeLEry, and SpaOTsc. Several methods that integrate scRNA-seq with ST data, including Cell2location [15], do not output inferred locations of cells. To compare inferred cell locations with ground truth location information, we used four metrics: Mean Squared Errors (MSE), Mean Absolute Errors (MAE), Average Jaccard Index, and Pearson’s Correlation. Detailed procedures on generating data, settings of running all methods, and evaluation metrics can be found in Supp. Info. Sec. 3.

Fig. 2b shows the comparison between TemSOMap and baseline methods using datasets simulated with a wide range of migration rates. For each migration rate, 20 simulated datasets were generated. We can observe that TemSOMap consistently outperforms other methods on all metrics and migration rates, indicating more accurate inferences of the cells’ spatial locations. From the accuracy changes of TemSOMap with migration rate, we observe that the performances tend to be worse for the migration rate at 0.5 and better when the migration rate is either small or large. This is because when the migration rate is low or high, the spatial distribution of cells will be more clustered based on either clones or cell types. Due to the partial consistency between clones and cell types [25], either pattern will benefit TemSOMap’s performance. When the migration rate is around 0.5, both cell type and clonal patterns are worse, making it more difficult for TemSOMap. However, TemSOMap is still able to outperform other methods under such circumstances. We provide some sample visualizations of the inferred spatial maps of cells under different migration rates (Supp. Fig. 2), which further shows the advantage of TemSOMap in inferring cells’ coordinates with similar clonal and cell-type patterns.

We also show the performances of TemSOMap compared with other methods on larger datasets (4096 cells, Supp. Fig. 3a) and with worse lineage barcode quality (25% dropouts, Supp. Fig. 3b). The comparisons are consistent overall with those on 1,024 cells datasets, even though the gap between TemSOMap and other methods decreases. This is because in both cases, the quality of the lineage barcodes is worse with dropouts, or with the same amount of mutations (the number of characters in the lineage barcode matrix) but with an increased sample size (1,024 cells to 4096 cells), which all caused the reconstructed cell lineage tree to be less accurate. We also compared the methods under different levels of batch effects in Supp. Fig. 3c. With the same migration rate, methods generally perform better with small batch effects compared to large batch effects, which is expected.

As mentioned in Methods, we use a default set of hyperparameters for these tests (specified in Supp. Info. Sec. 3.2). The number of clones used in the clonal loss is one of the hyperparameters, and here we specifically investigate the number of clones, as it can affect the effectiveness of the clonal loss. We conjecture that small number of clones allows robustness against the errors in the reconstructed cell lineage tree, so our default value is 4. Using simulated data, we proceed to compare TemSOMap’s performances under different numbers of clones (Supp. Fig. 3d). For the simulated datasets of 1,024 cells, we indeed observed that 4 clones yields the best overall performance. We recommend users to keep the number of clones small, and even 2 to 4 clones can still benefit the cell location inference as shown in the figure.

Using simulated datasets also allows us to perform an ablation test to quantitatively assess the contribution of different loss terms of TemSOMap. We started using only the expression similarity loss, then sequentially added the LA entropy loss, lineage loss, clonal loss, and variance loss. We can see that with the inclusion of each loss term, the performance of TemSOMap generally increases considering all four metrics (Fig. 2c).

Furthermore, we investigated the change in the values of the loss terms during optimization and found that under all different scenarios, training the total loss using the Adam optimizer, the different loss terms in TemSOMap converge simultaneously and reach a cell-spot mapping matrix that fits both the gene expressions and spatial patterns (Supp. Fig. 4a). Additionally, we compare the running time of TemSOMap with baseline methods (Supp. Fig. 4b). Compared with SpaOTsc, the other three methods scale better to large datasets, with CeLEry performing the fastest and TemSOMap being similar to Tangram. Detailed descriptions of the computational resources and other parameter settings for running all methods are in the Supp. Info. Sec. 3.4.

### 3.2 Testing TemSOMap on fitted mouse cortex data

In the previous section, we generated purely synthetic data to perform a comprehensive test of the methods with different migration rates (clonal patterns), dataset sizes, lineage barcode quality and batch effect sizes. Here we use a real ST dataset as a reference, to generate synthetic ST data with lineage barcodes that mimic the real ST dataset. The mammalian cerebral cortex is a suitable system to generate such datasets, as although there is no published real dataset with both spatial coordinates and lineage barcode information, its layered structure is well studied, and previous studies have shown the early development of the mammalian prefrontal cortex, with the birthtime of cells labeled at different embryonic times [16]. With this knowledge, we can simulate lineage barcode data for a real ST dataset on the cortex under the model of cell growth from inner to outer layers. We use the STARmap mouse brain cortex data from the Giotto toolbox [31, 9] with labeled cell types as the ST reference dataset (Fig. 3a). We use SpaTedSim to generate lineage barcodes and to fit the real STARmap dataset, where the spatial distribution of cell types is preserved.

**Fig. 3.**
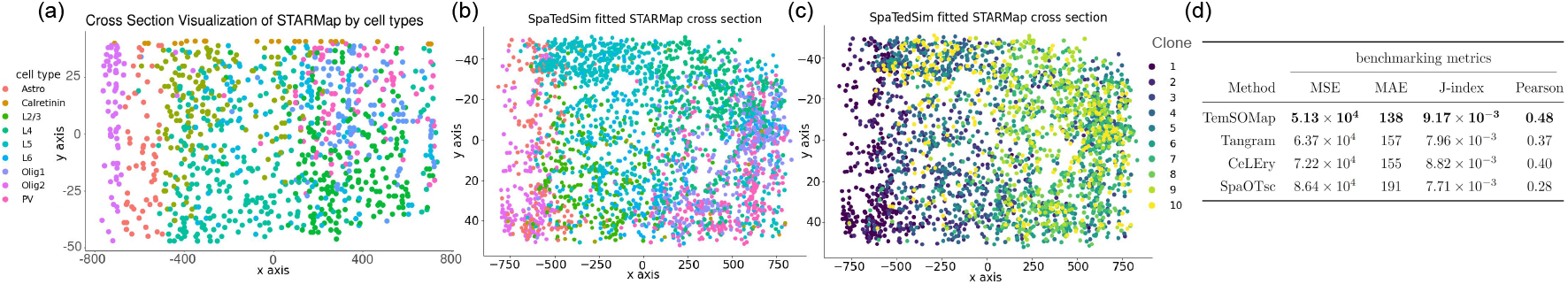
TemSOMap results on SpaTedSim-fitted STARmap mouse cortex data and comparison with other methods. (a) 2-D spatial visualization of a slice in the STARmap dataset. x,y axis show spatial coordinates(*µm*). (b) 2-D visualization of SpaTedSim-fitted STARmap data, colored by cell types. x,y axis show spatial coordinates(*µm*). (c) 2-D visualization of SpaTedSim-fitted STARMap data, colored by clones. x,y axis show spatial coordinates(*µm*). (d) Accuracy metrics of inferred single-cell coordinates by TemSOMap and the other state-of-the-art methods. Detailed descriptions of the methods and metrics are in Supp. Info. Sec. 3.

From the 3-D mouse cortex dataset, we first obtain a slice of 2-D distribution of cells by taking a cross section along the layers of the tissue (Supp. Fig. 5a). On the 2-D slice, we first separate spatial locations into clones based on the layered structure, and obtained ten clones [16] (Supp. Fig. 5b). For the area corresponding to each clone, we use a moving window to perform cell type density estimation to simulate realistic spatial distribution. Then, we merge the clones to get the final fitted data. The simulated dataset also aims to preserve other features of the reference STARmap data, such as cell type percentages (Supp. Fig. 5c) and average counts per cell type (Supp. Fig. 5d). Fig. 3b-c shows the visualization of generated data in 2-D space. Finally, we generated spot-level ST data with a fixed size (procedure in Supp. Info. Sec. 3.1) from the simulated STARmap data which is at single-cell resolution (Supp. Fig. 5e).

We used TemSOMap and other state-of-the-art methods to recover the hidden spatial coordinates of single cells by mapping the single-cell gene expressions onto the spot-level spatial data. We compared the performances of the methods using the same four metrics: MSE, MAE, Average Jaccard index, and Pearson’s correlation between the inferred cell coordinates and the ground truth. TemSOMap consistently outperformed the other methods and more accurately inferred the spatial coordinates of single cells (Fig. 3d).

### 3.3 Evaluating TemSOMap on E9.5 mouse embryo datasets

Understanding the spatiotemporal dynamics of early mammalian embryogenesis — that is, how a single omnipotent fertilized egg divides and differentiates into an embryo — is a fundamental problem in developmental biology. Integrating lineage tracing data and spatial transcriptomic data has the potential to uncover how cell types emerge and migrate in time and space. In this study, we attempt to integrate a lineage tracing dataset and a ST dataset of the mouse E9.5 embryo using TemSOMap. The lineage tracing dataset consists of paired scRNA-seq and CRISPR/Cas9-induced lineage barcodes of E9.5 mouse embryo cells [5]. The lineage barcodes are used to calculate the pairwise lineage distance and clonal labels of cells (Supp. Fig. 6a).

For the ST data, we used a Stereo-seq dataset [6] where spot-level ST data of an E9.5 mouse embryo with labeled cell types was provided. To map the single cells from the lineage tracing data onto the ST data, We first find the highly variable genes that are present in both datasets (based on the dispersion of genes, Supp. Fig. 6b). We then applied TemSOMap and obtained the mapping matrix *M*, which includes location information of cells in the lineage tracing dataset.

The Stereo-seq paper [6] provided cell type annotations of the high-resolution spots which correspond to different organs in the embryo (Fig. 4a, left). To visualize the lineage tracing data with cells in their inferred locations and compare with the ST data visualization in Fig. 4a, we need to annotate cells in the lineage tracing data with the same set of cell type labels. Therefore, we performed label transfer from the ST data to the lineage tracing data, by projecting the PCAs of lineage tracing data onto the PCA of the ST data, such that the two datasets are aligned in their gene expression space (Supp. Fig. 6c-d).

When obtaining inferred locations of cells from the *M* matrix, we have been using Eq. 11, which means a weighted mean location is used as the inferred location. An alternative way of determining inferred locations from *M* is to take the location with the highest probability for each cell, *i.e*., taking the Maximum A Posteriori (MAP) for each cell from the mapping matrix. To best explore this complex mouse embryo dataset, we used both modes, mean and MAP, as described above, to obtain inferred cell locations.

To compare with baseline methods, we also ran CeLEry and Tangram on this dataset. We did not run SpaOTsc, because it does not scale to this large dataset with 9707 cells in the lineage tracing dataset and 5031 spots in the ST dataset. For a fair comparison, we also used the two modes, mean and MAP to determine inferred cell locations for Tangram, as Tangram also outputs the *M* mapping matrix. Visualizations of the lineage tracing data with inferred locations using all methods and modes are shown in Supp. Fig. 7. We can see that results from CeLEry and Tangram-mean, though showing certain clustering patterns, preserve little shape of the embryo. TemSOMap-MAP and Tangram-MAP preserve the embryo and organ shapes best, and show overall comparable quality. Even though it is hard to quantify the performance differences between the two (due to the lack of ground truth on this real dataset), when comparing the results from TemSOMap-mean and Tangram-mean, we can see that TemSOMap-mean outperforms Tangram-mean significantly, indicating that the *M* matrix inferred by TemSOMap is likely more accurate than that of Tangram inference. Among TemSOMap-MAP and TemSOMap-mean, we chose to use the mapping results of TemSOMap-MAP for further analysis, as preserving the organ topology is critical for this dataset.

While better than other results, the organ patterns in TemSOMap-MAP results are still noisy. This can be due to multiple factors that pose difficulties for the mapping using TemSOMap. First, only a slice of the embryo is used in the reference ST data, while the lineage tracing dataset has all the cells in the embryo. Second, there can be a large batch effect between the two datasets, including individual differences from the two mouse embryos. However, when evaluating the spatial gene expression patterns of individual genes inferred by TemSOMap, we found that the TemSOMap-inferred spatial expression highly resembles the pattern observed in the ST data. Using inferred cell locations from TemSOMap, we can show spatial gene expression distributions of individual genes in the lineage tracing dataset. Furthermore, we can predict the spatial distribution of unseen genes or test genes. In Fig. 4b, the top panel shows examples of genes in the training set, while the bottom panels show examples of genes in the test set. For each gene, the left plot shows its expression level in the ST dataset, the middle plot shows its expression in the lineage tracing dataset, with inferred cell locations and the right plot shows the gene expression converted to spot resolution by calculating *M*^*T*^ *X*[∗, *j*], which results in a vector of length *n*_*spot*_ that represents the expression of gene *j* across spots. Details on performing this test and splitting the training and test sets are in Supp. Info. Sec. 3.3.

**Fig. 4.**
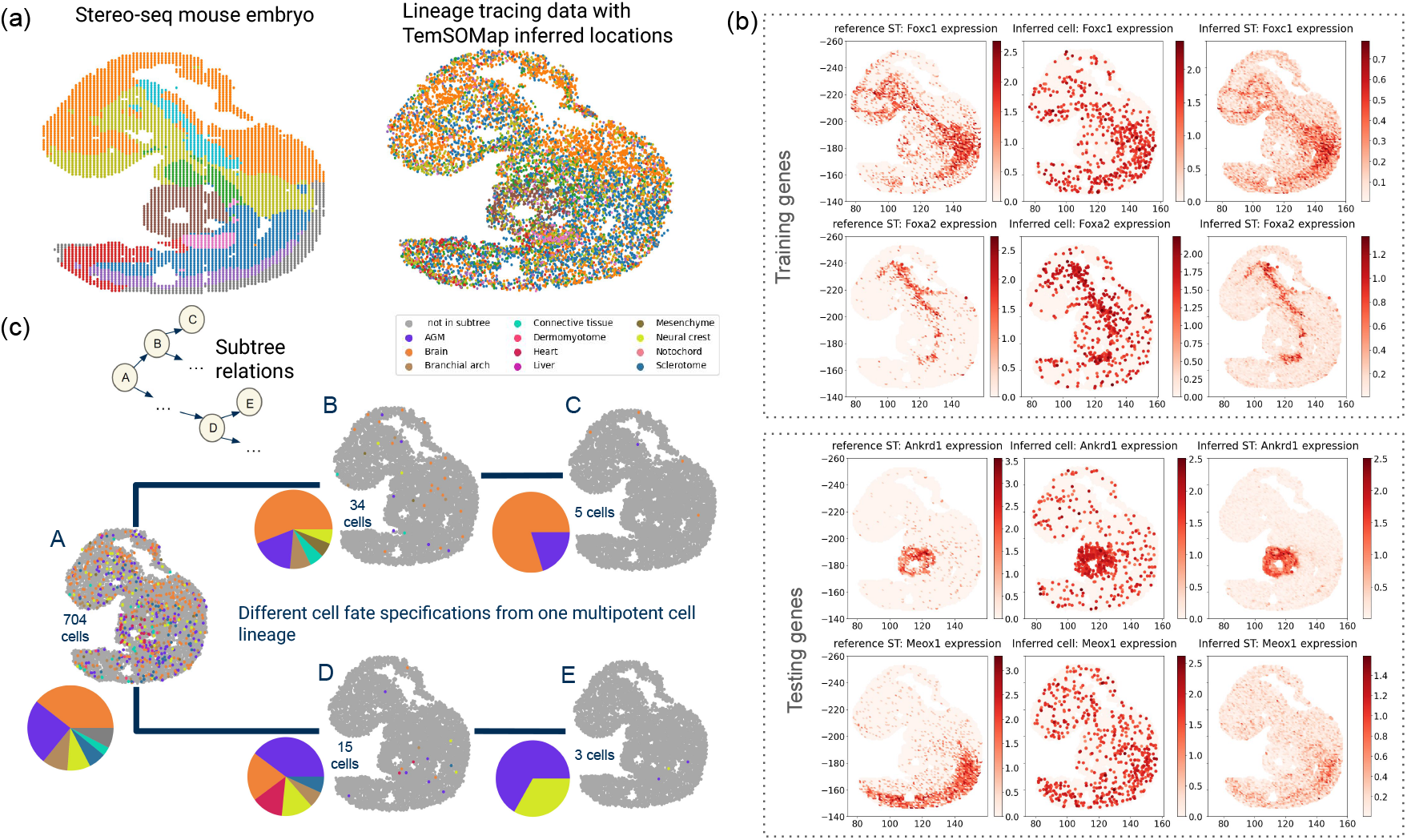
TemSOMap results on E9.5 mouse embryo data. (a) 2-D visualization of the input Stereo-seq mouse embryo data (spot level) and the TemSOMap-inferred mouse embryo data (single-cell level). Colors represent cell types from the Stereo-seq annotation. (b) Comparing spatial maps of gene expressions of the reference ST data (left), inferred cell-level data (middle), and inferred spot-level data (right). Foxc1 and Foxa2 are used in the training of TemSOMap; Ankrd1 and Meox1 are masked from the training of TemSOMap and their spatial distribution is predicted using TemSOMap. (c) Spatiotemporal analysis on the cell fate specifications on the cell lineage. Each embryo plot represents the spatial distribution of cells for each subtree, with pie plots showing the cell type percentages of the leaves.

More spatial gene comparisons are shown in Supp. Fig. 8 (for genes in the training set) and Supp. Fig. 9 (for genes in the test set). From the visualizations, we can see that for genes in both the training set and the test set, the TemSOMap predicted spatial distribution of gene expression in the lineage tracing dataset highly resembles that in the ST dataset, especially the inferred ST level data (the rightmost plots for each gene). These results not only support the inferred cell locations, but also indicate the potential of TemSOMap in compensating for the lack of gene throughput or resolution of ST technologies.

Finally, the lineage tracing dataset with inferred cell locations allows us to gain more insights into the spatiotemporal changes of cells’ locations along with their cell division histories (Fig. 4b). With mapping results from TemSOMap (Fig. 4a right), we analyze a subtree of the lineage tree, which corresponds to a large multipotent progenitor (node A in Fig. 4b). This progenitor is the common ancestor of 704 observed present-day cells, whose spatial locations and cell types can be visualized (Fig. 4b). Going down the lineage from this node on the lineage tree, we can observe that different daughter cells take on different cell fate specifications; while one lineage (*A* → *B* → *C*) differentiates into mostly brain cells, another lineage (*A* → *D* → *E*) differentiates into a majority of AGM cells.

### 3.4 Evaluating TemSOMap on baseMEMOIR mESC lineages

Moreover, recent advances in spatiotemporal lineage tracing technologies have enabled simultaneous readouts of lineage barcodes, gene expressions, and spatial coordinates of single cells. Although such technologies are still in their early development, with low throughput and sample size, they provide valuable test cases for cell location inference.

We use the recently developed dataset, baseMEMOIR, which provides lineage barcodes, gene expressions, and spatial coordinates derived from mouse embryonic stem cells (mESCs). The dataset contains multiple small colonies (10 - 50 cells) of 12 targeted gene expressions. The lineage barcodes can be used to calculate the pairwise cophenetic distances between cells. To set up the experiment for cell location inference. We mask the spatial coordinates, and add batch effects to the cell-by-gene matrix to produce scRNA-seq data. Then, we use TemSOMap to infer the spatial coordinates of cells and compare them with the ground truth. Moreover, we use the inferred lineage tree using the BEAST2 analysis from the original papers to obtain clonal IDs for the cells.

We show comparisons between the spatial map of cells on both ground truth coordinates and inferred coordinates using TemSOMap and the state-of-the-art method, Tangram. Labeled by both cell types and clonal IDs, we observe that TemSOMap can accurately infer the spatial coordinates of cells and maintain similar cell types and clonal clusters. By calculating Mean-Squared Errors(MSE), it show that TemSOMap outperforms Tangram consistently on all colonies.

### 3.5 Evaluating TemSOMap on baseMEMOIR mESC lineages

## 4 Discussion

We presented TemSOMap, a method that can incorporate cell lineage (temporal) information when mapping scRNA-seq data to ST data. This not only allows for more accurate inference of cell spatial locations in scRNA-seq data, but also generates integrated lineage-resolved spatial data; that is, a dataset that has three modalities: gene expression, lineage and clonal information, and (inferred) spatial locations of cells. We have shown that such spatiotemporal data can be used to study cell fate and cell location specifications during tissue development. Although TemSOMap takes advantage of the assumption that cells from the same clone tend to located closely in space, our simulation tool SpaTedSim allows us to generate data with different migration rates, which is the probability for a cell to migrate from its sister cell. We can see that TemSOMap still outperforms baseline methods even when the migration rate is high. This means that TemSOMap exploits clonal and lineage patterns even when such signals are very noisy.

In this paper, we primarily used lineage tracing data to obtain cell lineage and clonal information. Other types of data can potentially be used to obtain this information. For example, literature has suggested that scATAC-seq data or scRNA-seq data can include information on mitochondria somatic mutations, which can be used to trace cell lineages [22, 17, 23]. Moreover, for cancer tissues, copy number variation (CNV) that can be detected from scRNA-seq or scATAC-seq data can also be used to trace cell lineages [22]. TemSOMap is applicable to this wide variety of data types.

TemSOMap can also be applied to datasets from a wide range of ST technologies. In this paper, we have used STARmap data and Steoro-seq data, and the simulated datasets were designed to simulate larger spatial spots like those in 10x Visium (around 50 *µm* [1]). The Steoro-seq data has subcellular resolution, though we have used the pre-processed data in the Steoro-seq paper where high-resolution spots are binned into larger spots. But TemSOMap can also work with subcellular resolution data such as Xenium(0.5*µm* [14]) since the mapping matrix in TemSOMap can be implied as a subcellular-level mapping of cells’ transcripts from cells to space.

With the continuing development of both ST and lineage tracing technologies, spatio-temporal modeling is increasingly important and feasible to uncover biological insights. Emerging technologies that sequence spatial and temporal information simultaneously [13] can be used to test or train new spatio-temporal models. We anticipate that pioneering efforts like TemSOMap will drive the development of future spatio- temporal methods to further harness the potential of these cutting-edge technologies.

## References

1. 10x Genomics. Spatial gene expression. https://www.10xgenomics.com/products/spatial-gene-expression.

2. Lyla Atta and Jean Fan. Computational challenges and opportunities in spatially resolved transcriptomic data analysis. Nat. Commun., 12(1):5283, September 2021.

3. Tommaso Biancalani, Gabriele Scalia, Lorenzo Buffoni, Raghav Avasthi, et al. Deep learning and alignment of spatially resolved single-cell transcriptomes with tangram. Nature Methods, 18(11):1352–1362, October 2021.

4. Zixuan Cang and Qing Nie. Inferring spatial and signaling relationships between cells from single cell transcriptomic data. Nat. Commun., 11(1):2084, April 2020.

5. Michelle M. Chan, Zachary D. Smith, Stefanie Grosswendt, Helene Kretzmer, et al. Molecular recording of mammalian embryogenesis. Nature, 570(7759):77–82, Jun 2019.

6. Ao Chen, Sha Liao, Mengnan Cheng, Kailong Ma, et al. Spatiotemporal transcriptomic atlas of mouse organogenesis using dna nanoball-patterned arrays. Cell, 185(10):1777– 1792.e21, May 2022.

7. Kok Hao Chen, Alistair N Boettiger, Jeffrey R Moffitt, Siyuan Wang, et al. RNA imaging. spatially resolved, highly multiplexed RNA profiling in single cells. Science, 348(6233):aaa6090, April 2015.

8. Ruben Dries, Jiaji Chen, Natalie Del Rossi, Mohammed Muzamil Khan, et al. Advances in spatial transcriptomic data analysis. Genome Res., 31(10):1706–1718, October 2021.

9. Ruben Dries, Qian Zhu, Rui Dong, Chee-Huat Linus Eng, et al. Giotto: a toolbox for integrative analysis and visualization of spatial expression data. Genome Biology, 22(1), March 2021.

10. Wuming Gong, Hyunwoo J Kim, Daniel J Garry, and Il-Youp Kwak. Single cell lineage reconstruction using distance-based algorithms and the R package, DCLEAR. BMC Bioinformatics, 23(1):103, March 2022.

11. Shanshan He, Ruchir Bhatt, Carl Brown, Emily A Brown, et al. High-plex multiomic analysis in FFPE at subcellular level by spatial molecular imaging. bioRxiv, page 2021.11.03.467020, November 2021.

12. Matthew G Jones, Alex Khodaverdian, Jeffrey J Quinn, Michelle M Chan, Jeffrey A Hussmann, Robert Wang, Chenling Xu, Jonathan S Weissman, and Nir Yosef. Inference of single-cell phylogenies from lineage tracing data using cassiopeia. Genome Biology, 21(1), April 2020.

13. Matthew G. Jones, Dawei Sun, Kyung Hoi (Joseph) Min, William N. Colgan, et al. Spatiotemporal lineage tracing reveals the dynamic spatial architecture of tumor growth and metastasis. October 2024.

14. Rongqin Ke, Marco Mignardi, Alexandra Pacureanu, Jessica Svedlund, et al. In situ sequencing for RNA analysis in preserved tissue and cells. Nat. Methods, 10(9):857–860, September 2013.

15. Vitalii Kleshchevnikov, Artem Shmatko, Emma Dann, Alexander Aivazidis, et al. Cell2location maps fine-grained cell types in spatial transcriptomics. Nat. Biotechnol., January 2022.

16. Sharon M. Kolk and Pasko Rakic. Development of prefrontal cortex. Neuropsychopharmacology, 47(1):41–57, October 2021.

17. Caleb A. Lareau, Vincent Liu, Christoph Muus, et al. Mitochondrial single-cell atac-seq for high-throughput multi-omic detection of mitochondrial genotypes and chromatin accessibility. Nature Protocols, 18(5):1416–1440, February 2023.

18. Je Hyuk Lee, Evan R Daugharthy, Jonathan Scheiman, Reza Kalhor, et al. Fluorescent in situ sequencing (FISSEQ) of RNA for gene expression profiling in intact cells and tissues. Nat. Protoc., 10(3):442–458, March 2015.

19. Bin Li, Wen Zhang, Chuang Guo, Hao Xu, et al. Benchmarking spatial and single-cell transcriptomics integration methods for transcript distribution prediction and cell type deconvolution. Nat. Methods, May 2022.

20. Chang Liu, Rui Li, Young Li, Xiumei Lin, et al. Spatiotemporal mapping of gene expression landscapes and developmental trajectories during zebrafish embryogenesis. Dev. Cell, 57(10):1284–1298.e5, May 2022.

21. Sophia K Longo, Margaret G Guo, Andrew L Ji, and Paul A Khavari. Integrating single-cell and spatial transcriptomics to elucidate intercellular tissue dynamics. Nat. Rev. Genet., 22(10):627–644, October 2021.

22. Leif S Ludwig, Caleb A Lareau, Jacob C Ulirsch, Elena Christian, et al. Lineage tracing in humans enabled by mitochondrial mutations and single-cell genomics. Cell, 176(6):1325–1339.e22, March 2019.

23. Lena Nitsch, Caleb A Lareau, and Leif S Ludwig. Mitochondrial genetics through the lens of single-cell multiomics. Nat. Genet., 56(7):1355–1365, July 2024.

24. Xinhai Pan, Hechen Li, Pranav Putta, and Xiuwei Zhang. LinRace: cell division history reconstruction of single cells using paired lineage barcode and gene expression data. Nat. Commun., 14(1):8388, December 2023.

25. Xinhai Pan, Hechen Li, and Xiuwei Zhang. Tedsim: temporal dynamics simulation of single-cell rna sequencing data and cell division history. Nucleic Acids Research, 50(8):4272–4288, April 2022.

26. Bushra Raj, Daniel E. Wagner, Aaron McKenna, Shristi Pandey, et al. Simultaneous single-cell profiling of lineages and cell types in the vertebrate brain. Nature Biotechnology, 36(5):442–450, May 2018.

27. N Saitou and M Nei. The neighbor-joining method: a new method for reconstructing phylogenetic trees. Mol. Biol. Evol., 4(4):406–425, July 1987.

28. Irepan Salvador-Martínez, Marco Grillo, Michalis Averof, and Maximilian J Telford. Is it possible to reconstruct an accurate cell lineage using crispr recorders? eLife, 8:e40292, jan 2019.

29. Bastiaan Spanjaard, Bo Hu, Nina Mitic, Pedro Olivares-Chauvet, et al. Simultaneous lineage tracing and celltype identification using crispr–cas9-induced genetic scars. Nature Biotechnology, 36(5):469–473, May 2018.

30. Michael Spencer Chapman, Anna Maria Ranzoni, Brynelle Myers, Nicholas Williams, et al. Lineage tracing of human development through somatic mutations. Nature, 595(7865):85–90, May 2021.

31. Xiao Wang, William E Allen, Matthew A Wright, Emily L Sylwestrak, et al. Three-dimensional intact-tissue sequencing of single-cell transcriptional states. Science, 361(6400), July 2018.

32. Liangchen Yue, Feng Liu, Jiongsong Hu, Pin Yang, et al. A guidebook of spatial transcriptomic technologies, data resources and analysis approaches. Comput. Struct. Biotechnol. J., 21:940–955, January 2023.

33. Qihuang Zhang, Shunzhou Jiang, Amelia Schroeder, Jian Hu, et al. Leveraging spatial transcriptomics data to recover cell locations in single-cell RNA-seq with CeLEry. Nat. Commun., 14(1):4050, July 2023.

