## Supplementary Information for "Mapping lineage-resolved scRNA-seq data with spatial transcriptomics using TemSOMap"

#### 1 Supplementary Figures

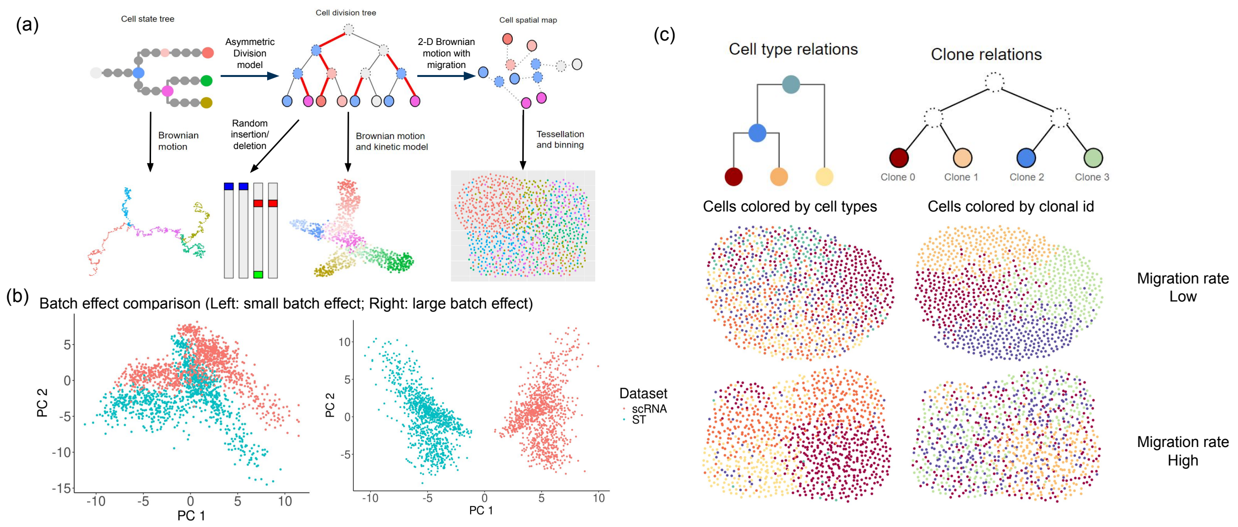


Supplementary Figure 1: (a) SpaTedSim simulation framework. SpaTedSim takes the input of a cell state tree, a cell and a cell division tree, and outputs paired lineage barcodes, gene expressions and spatial coordinates. (b) Batch effect visualization of SpaTedSim generated gene expression. The ST batch represents the single-cell gene expressions before the tessellation of the space into spots. (c) Visualizing the cell type/clonal pattern of SpaTedSim simulated datasets. The two trees are the ground truth cell state tree and division tree input to SpaTedSim. With a lower migration rate, the clonal patterns are clearer; with a higher migration rate, cell type patterns are clearer.

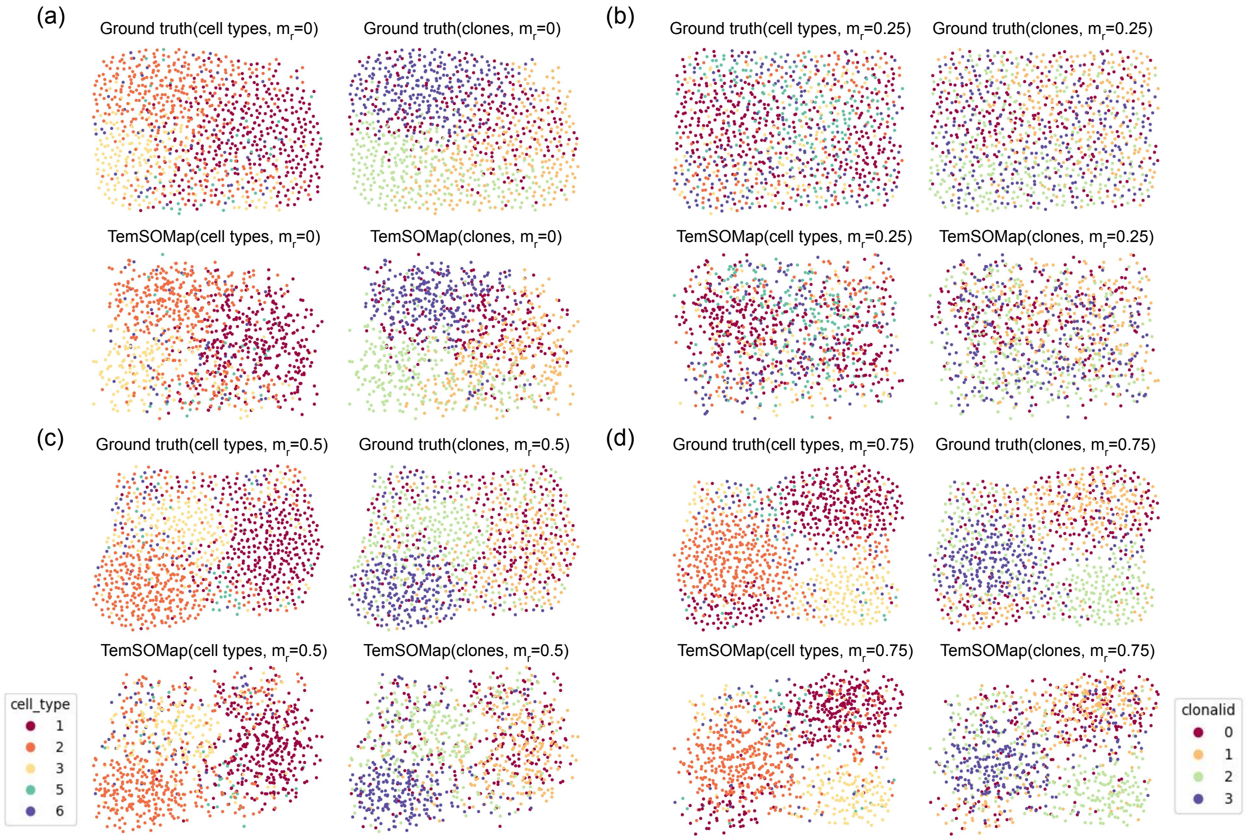


Supplementary Figure 2: Visualizing TemSOMap inferred spatial map of cells comparing with the ground truth. With varying migration rates, the spatial patterns of the simulated data change. The left column in each subfigure shows the cell type pattern, and the right column shows the clonal pattern. (a) migration rate is 0, means no migration happens. (b) migration rate is 0.25. (c) migration rate is 0.5. (d) migration rate is 0.75.

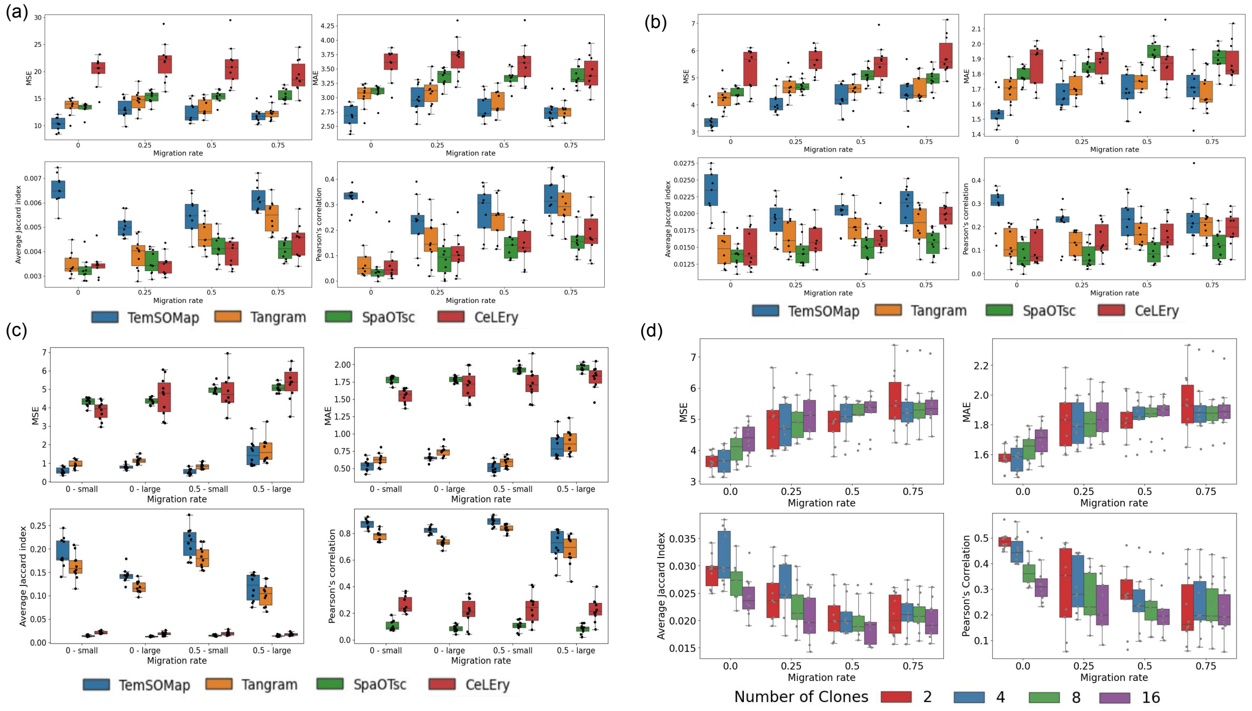


Supplementary Figure 3: (a) Comparisons of TemSOMap, and other baseline methods on SpaTedSim-simulated datasets, with varying migration rates, on 4096 cells for the datasets. For MSE and MAE, lower values mean better accuracy. For the Average Jaccard Index and Pearson's Correlation, higher values mean better accuracy. (b) Comparisons of TemSOMap with worse input lineage barcodes, and other baseline methods on SpaTedSim-simulated datasets. Dropouts are introduced by randomly dropping 25% of the lineage barcodes, which lower the quality of the input to TemSOMap while the other methods perform the same. This test shows TemSOMap's performance despite the dropouts in the lineage data. (c) Comparisons of TemSOMap, and other baseline methods on SpaTedSim-simulated datasets, with varying migration rates and batch effects. Each group of boxes corresponds to a combination of migration rate and batch effect size. For example, "0.5-small" means data generated with migration rate 0.5 and small batch effect. Detailed descriptions of the batch effects can be found in Supp. Info. Sec. 2.2. (d) Comparisons of TemSOMap performances with varying numbers of clones. The clonal ids for single cells are generated by cutting the rebalanced cell lineage.

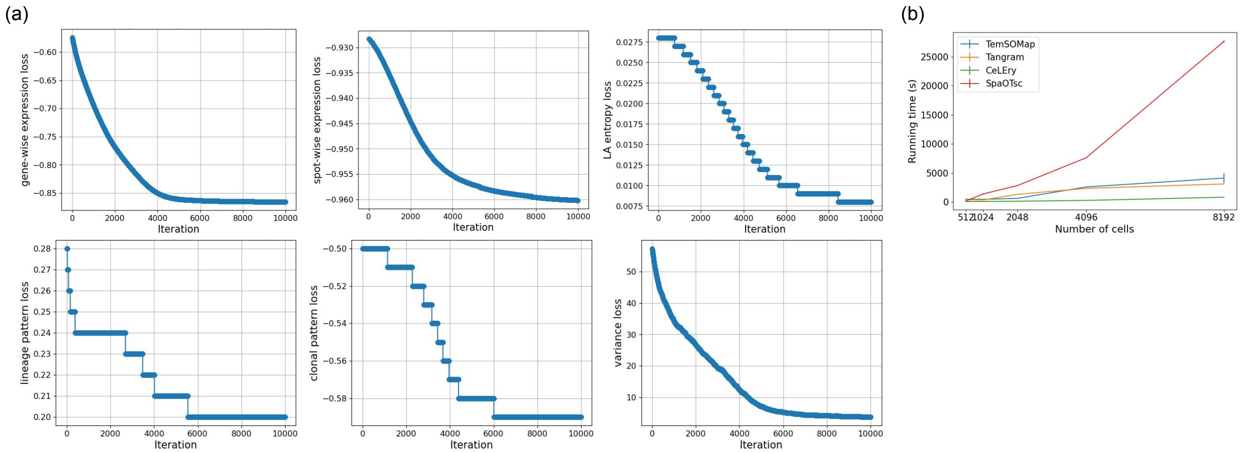


Supplementary Figure 4: (a) Loss term curves with the increase of iterations. From left to right are the spot-wise expression loss, gene-wise expression loss, location-aware entropy loss, lineage loss, clonal loss, and variance loss. All loss terms converge together with the optimization of the total loss using Adam optimizer. (b) Running time comparisons for the four methods. With a fixed spot size, we increase the number of cells which accordingly increases the number of spots (Approximately $n\_spots = \log_2(n\_cells)$). For each dataset size, we record the average running time across five datasets. More details are in Supp. Info. Sec. 3.1.

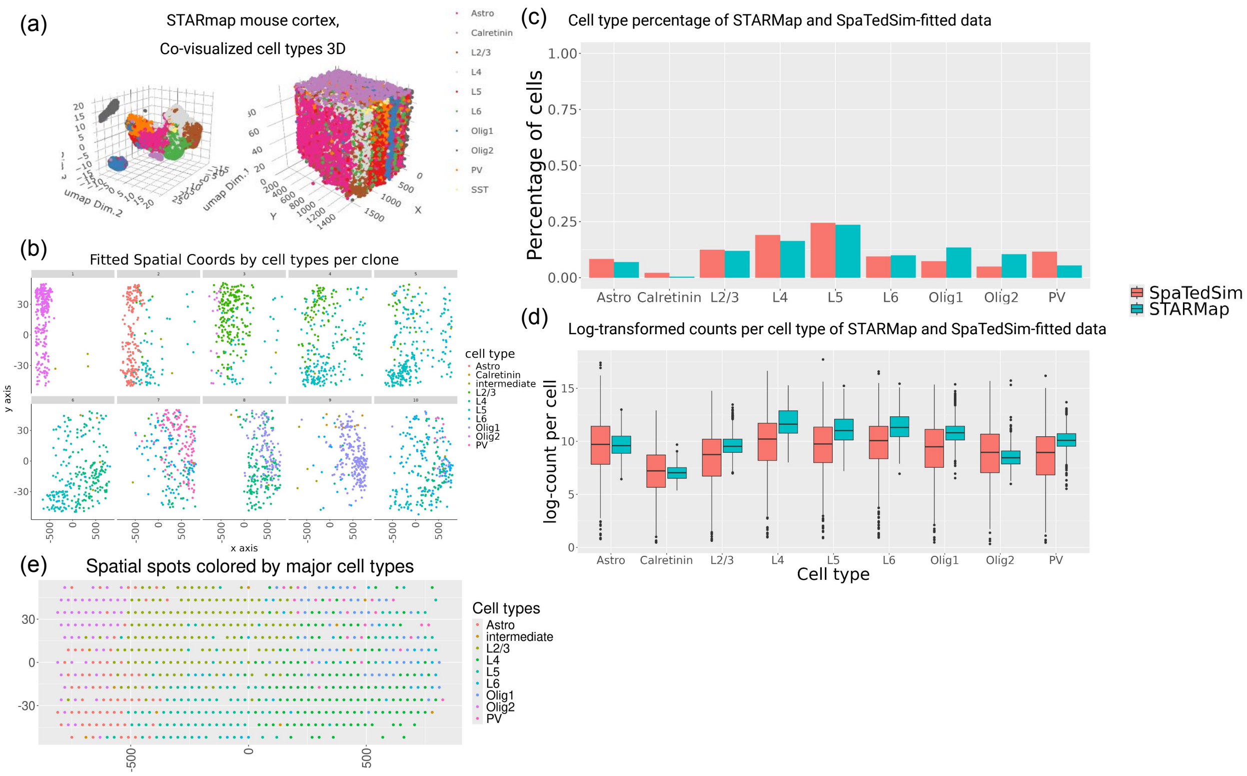


Supplementary Figure 5: (a) 3-D visualization of the STARmap mouse cortex data, colored by cell types. The left panel is the 3-D UMAP visualization of the gene expressions, and the right panel is the 3-D spatial visualization of the mouse cortex. (b) 2-D visualization of the spatial coordinates of SpaTedSim-fitted mouse cortex data, by clones. Each clone is generated using a moving window to subset the real dataset for cell type density estimation. SpaTedSim also introduces intermediate cell types based on the cell state tree. (c) A comparison of cell type percentages by cell types, comparing SpaTedSim-fitted data(2560 cells, 10 clones) and the 2-D cross section of the STARmap data (1046 cells). (d) A comparison of log-transformed gene counts of cells, comparing SpaTedSim-fitted data and the 2-D cross section of the STARmap data, grouped by cell types. (e) Visualization of the output spot-level coordinates, colored by major cell types at each spot. The expression at the spots are calculated by summing up the counts of the cells within the tessellation area of the spots.

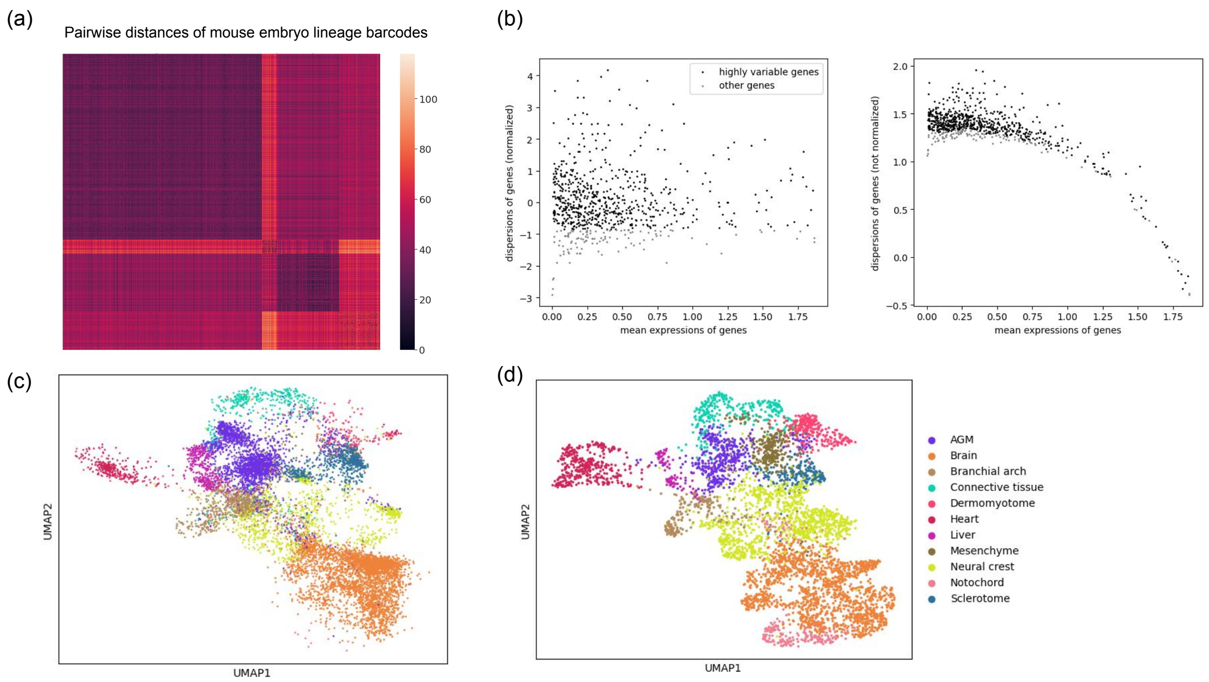


Supplementary Figure 6: (a) Heatmap of the pairwise hamming distance matrix of mouse embryo lineage barcodes. The lineage barcode data is from the embryo 2 in [1]. (b) Filtering highly variable genes of the expression data. Top 400 highly variable genes that are intersection between the spatial and lineage dataset are used to perform TemSOMap mapping, and 193 other highly variable genes are used to test the prediction of TemSOMap. (c) UMAP visualization of the lineage data, colored by cell types annotation transferred from the stereo-seq data. (d) UMAP visualization of the stereo-seq data, colored by cell types annotation from the Giotto toolbox [2].

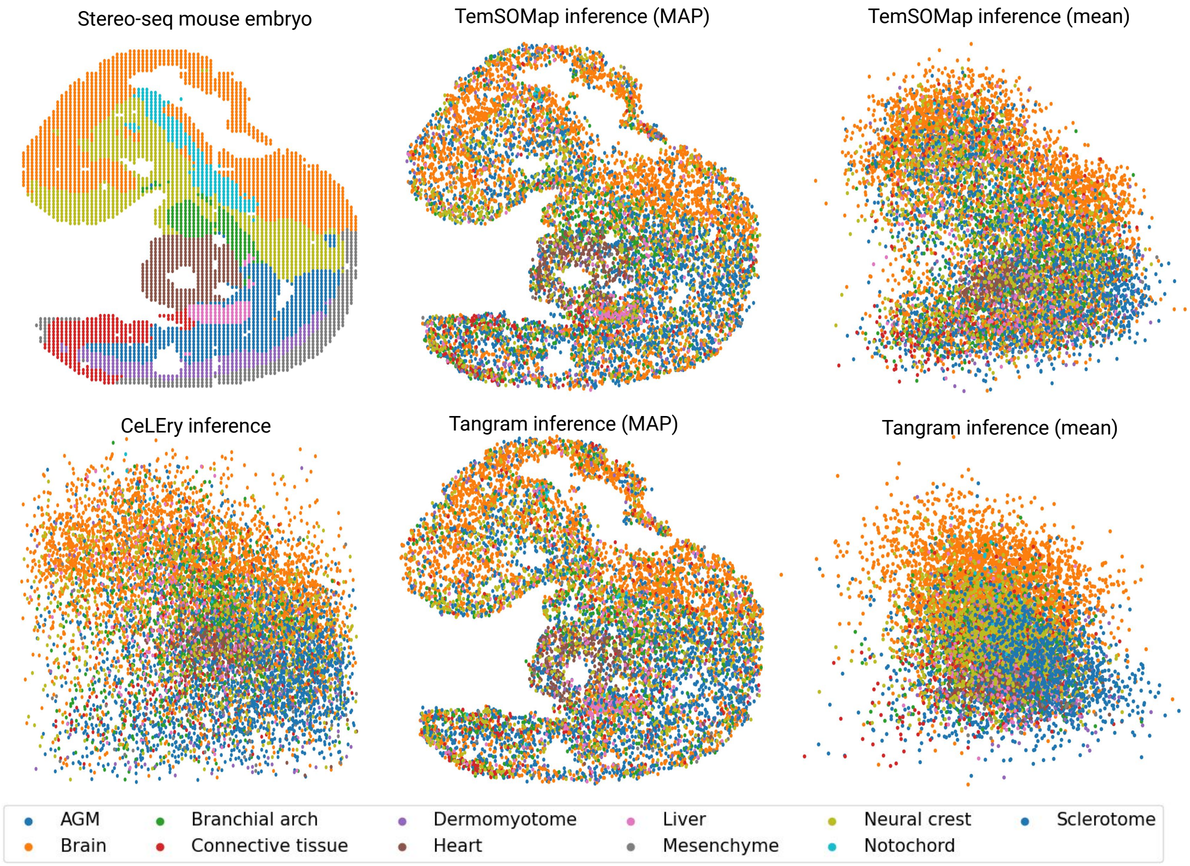


Supplementary Figure 7: Visualizations of the spatial map of the E9.5 mouse embryo, comparing the reference stereo-seq data, TemSOMap inferred data using both MAP and mean predicted coordinates, CeLEry inferred data and Tangram inferred data using both MAP and mean predicted coordinates.

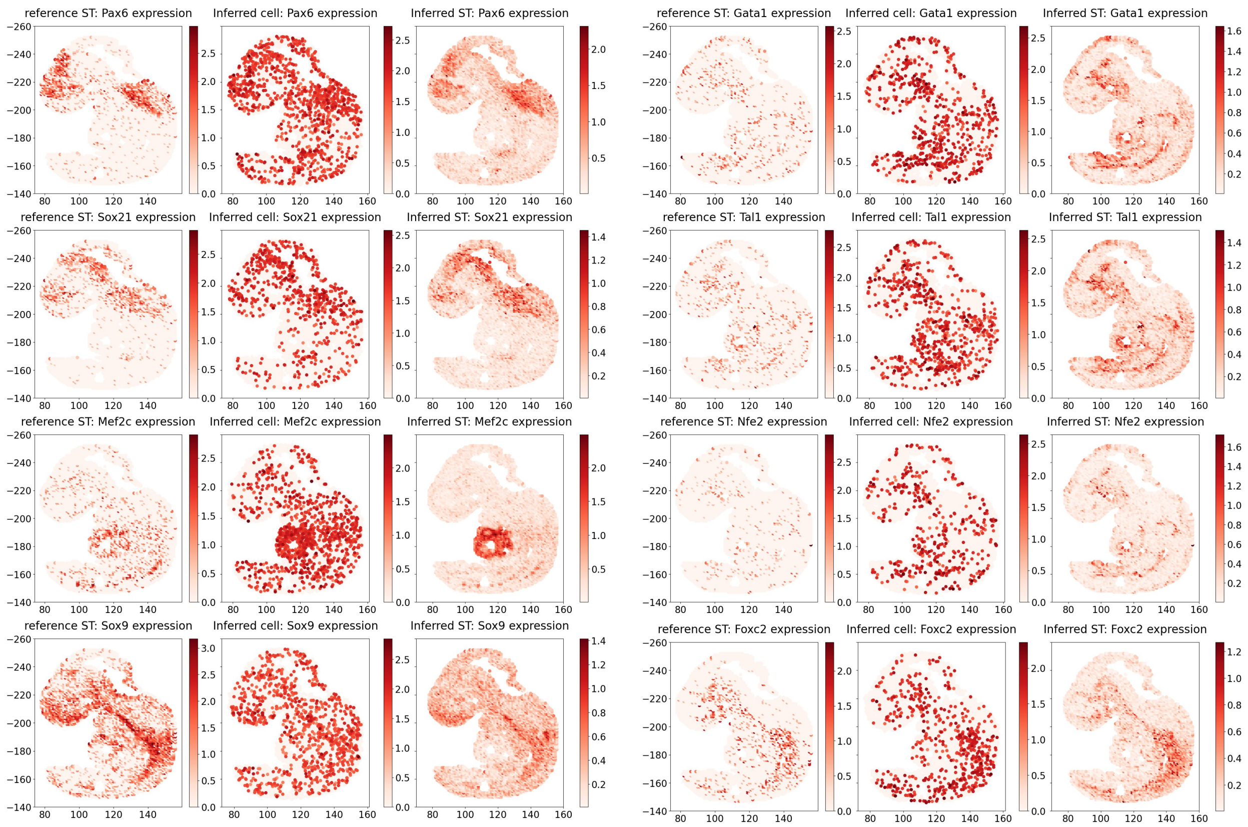


Supplementary Figure 8: Marker Spatial genes visualization of mapped single-cell mouse embryo data (Training set). These genes are used in the training process of TemSOMap. The three columns compare the spatial maps of spatial gene expressions of the reference ST data, inferred cell-level data, and inferred spot-level data.

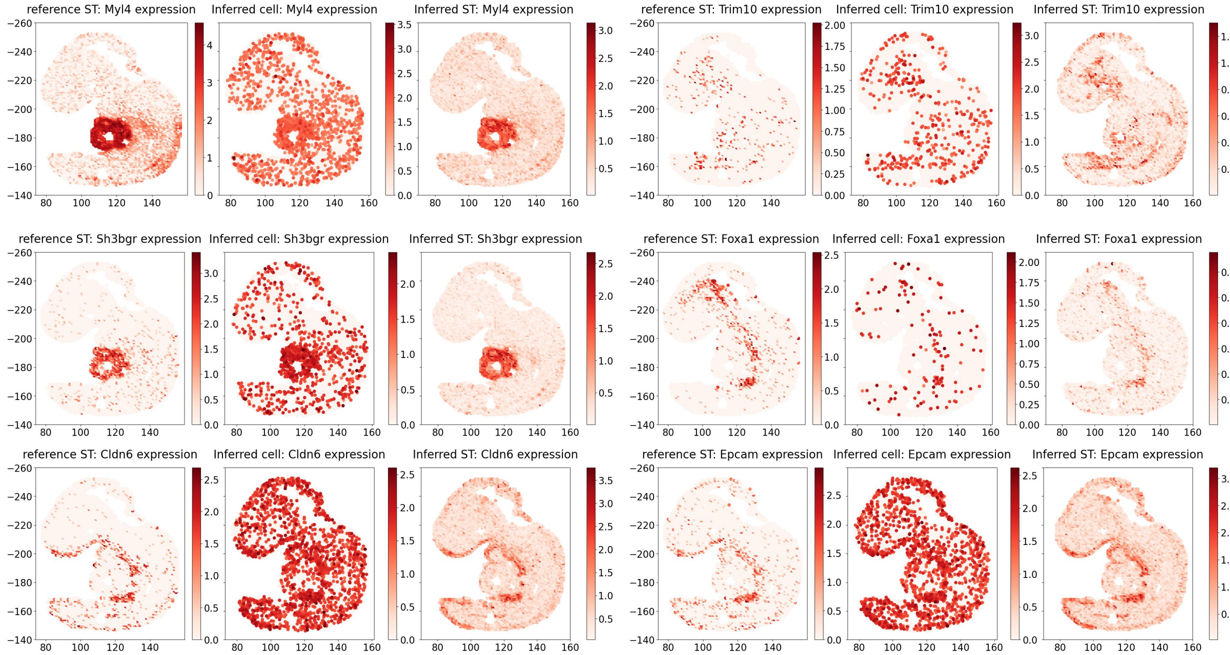


Supplementary Figure 9: Marker Spatial genes visualization of mapped single-cell mouse embryo data (Testing set). These genes are masked from the training process of TemSOMap. The three columns compare the spatial maps of spatial gene expressions of the reference ST data, inferred cell-level data, and inferred spot-level data.

### 2 Simulating paired gene expression, lineage barcodes and spatial coordinates using SpaTedSim

To characterize the spatio-temporal dynamics of cells' transcriptomes, and provide a tool to test computational methods for multimodal data integration, we developed SpaTedSim, a spatio-temporal dynamics simulation of single cells. SpaTedSim generates paired single-cell RNA counts, spatial coordinates, and lineage barcodes simultaneously, by simulating realistic biological events including cell symmetric and asymmetric divisions, and cell migrations. SpaTedSim can tune the parameters for cell division and migration events that lead to changes in the spatio-temporal correlation of cells' gene expressions, accommodating various biological circumstances. SpaTedSim can test different types of computational tasks, including trajectory inference, lineage inference, spatial transcriptome mapping, etc. SpaTedSim can also fit real ST data and generate synthetic datasets that share realistic spatial gene expression with clonal information.

#### 2.1 Simulating cells' lineage barcode and gene expressions on the cell division tree

we adopt the TedSim framework: We start with two tree structures: a cell state tree, indicating the developmental trajectory of cell types; and a cell division tree, indicating the cell division history. TedSim first runs a Brownian motion model on the cell state tree to obtain the mean latent space representations for each cell state. TedSim then runs another Brownian motion model on the cell division tree to obtain the latent space representations and then transform the latent space into gene expression levels for each cell. A cell's latent space representation can be regarded as a weighted sum of the state mean representations and the cell's lineage representations. We adopt the asymmetric division model to determine the cell states of all cells while performing Brownian motion on the lineage tree. Starting from the root of the cell lineage tree, a cell can divide either symmetrically into two cells of the same state as their parent or asymmetrically, which means one child cell stays at the same state and the other child cell shifts to a future state on the cell state tree. The leaf cells on the cell division tree are considered present-day cells, and the gene expression profiles of the leaf cells are the output transcriptomic data for a scRNA-seq experiment. A detailed description of the simulation of gene expression and lineage barcode can be found in TedSim [3].

#### 2.2 Adding batch effect between scRNA-seq data and spatial transcriptomic data

It is universally acknowledged that scRNA-seq experiments conducted in different laboratories and at different times contain batch effects that may compromise the integration and interpretation of the data. We implemented a function to add different levels of batch effects to simulated cells' gene expressions. This function can help to test how TemSOMap and other state-of-the-art methods handle batch effects between the reference ST data and the query scRNA-seq data. The Pseudocode to add batch effect between scRNA-seq and ST data is given below:

– **Input:**
  - $counts$: RNA-seq counts matrix (true counts, before batch effect)
  - $ncells$: Number of cells (for each batch)
  - $ngenes$: Number of genes
  - $batch\_effect\_size$: Size of the batch effect (default = 1)
– **Procedure:**
  a) Create a vector $batchIDs$ that assigns a batch label to each cell:
     - Assign batch 1 to the first $ncells$ cells, indicating the scRNA-seq data.
     - Assign batch 2 to the next $ncells$ cells, indicating the ST data before binning.
  b) Initialize a matrix $mean\_matrix$ of size $ngenes \times 2$ to store batch-specific mean values for each gene.
  c) Generate a random vector $gene\_mean$ of length $ngenes$ from a normal distribution with mean 0 and standard deviation 1.
  d) For each gene $igene$ from 1 to $ngenes$:

(a) For each batch (1 and 2), generate a random value between $gene\_mean[igene] - batch\_effect\_size$ and $gene\_mean[igene] + batch\_effect\_size$.
(b) Store the two values in the corresponding row of $mean\_matrix$.

e) Initialize a matrix $batch\_factor$ of size $(2 \times ncells) \times ngenes$ to store the batch effect for each cell and gene.

f) For each gene $igene$ from 1 to $ngenes$:
(a) For each cell $icell$ from 1 to $2 \times ncells$:
- Sample a random value from a normal distribution with mean based on gene and batchID: $mean\_matrix[igene, batchIDs[icell]]$ and standard deviation 0.01.
- Store this value in $batch\_factor[icell, igene]$.

g) Apply the batch effect to the RNA-seq counts:
- Compute $counts\_after$:

$$counts\_after \leftarrow \text{round}\left(2^{\log_2(\text{counts})+\text{batch\_factor}[1:ncells,]}\right)$$

- Compute $observed\_counts\_matrix\_after$:

$$observed\_counts\_matrix\_after \leftarrow \text{round}\left(2^{\log_2(\text{counts})+\text{batch\_factor}[ncells+1:2\times ncells,]}\right)$$

– **Output:**
- $counts\_after$: Counts matrix for the first batch after applying the batch effect.
- $observed\_counts\_matrix\_after$: Counts matrix for the second batch after applying the batch effect.

##### 2.3 Post-processing of simulated spatial coordinates of cells

Post-processing has two steps to make the overall spatial distribution of cells more uniform: 1. Pull the outlier cells that are too far away from the neighbors; 2. Push the cells too close to each other based on a given minimum distance. Here are the pseudocodes that describes the two steps:

###### Pseudocode for pulling cells based on maximum distance constraint

– **Input:**
- spatial coordinates of cells (x, y)
- $max\_distance$: Maximum allowed distance between points
- $k$: The $k$-th closest point to check (default is 3)
- $iterations$: Number of iterations to perform (default is 100)
- $step\_size$: Step size for adjusting points (default is 0.05)

– **Procedure:**

a) For each iteration from 1 to $iterations$:
(a) Recalculate the distance matrix $D$ between all cells $D = \{D_{ij} = ||(x_i - x_j)^2 + (y_i - y_j)^2||_2\}$
(b) For each cell $(x_j, y_j)$:
i. Find the $k$-th closest cell to cell $j$ in $D_{*,j}$
ii. If the distance to the $k$-th closest cell exceeds $max\_distance$:
A. Identify the index $l$ of the $k$-th closest cell
B. Compute the vector $\vec{v}$ from cell $j$ to cell $l$
C. Normalize $\vec{v}$ and scale it by $step\_size$
D. Adjust cell $j$'s spatial coordinates by subtracting the components of the scaled vector: $(x'_j, y'_j) = (x_j, y_j) - step\_size \cdot \frac{\vec{v}}{|\vec{v}|}$
(c) Recalculate the distance matrix after the adjustment

b) If the distance of all points to their $k$-th closest neighbor is less than or equal to $max\_distance$, break the loop and return the updated spatial coordinates of cells

– **Output:**
- Adjusted spatial coordinates of cells $(x', y')$.

**Pseudocode for pushing cells based on minimum distance constraint**

- **Input:**
  - Spatial coordinates of cells (x, y)
  - *min_distance*: Minimum allowed distance between points
  - *iterations*: Maximum number of iterations to perform (default is 20)
  - *step_size*: Step size for adjusting points (default is 0.05)
- **Procedure:**
  a) For each iteration from 1 to *iterations*:
     (a) Recompute the distance matrix between the spatial coordinates of all cells: $D = \{D_{ij} = ||(x_i - x_j)^2 + (y_i - y_j)^2||_2\}$
     (b) For each cell $(x_j, y_j)$:
        i. Identify all cells closer to $j$ than *min_distance* but not at a zero distance
        ii. For each point $k$ that is too close:
           A. Compute the vector $\vec{v}$ from point $j$ to point $k$
           B. Normalize $\vec{v}$ and scale it by *step_size*
           C. Adjust spatial coordinates of cell $j$ by adding the components of the scaled vector $(x'_j, y'_j) = (x_j, y_j) + step\_size \cdot \frac{\vec{v}}{|\vec{v}|}$
           D. Adjust spatial coordinates of cell $k$ by subtracting the components of the scaled vector: $(x'_k, y'_k) = (x_k, y_k) - step\_size \cdot \frac{\vec{v}}{|\vec{v}|}$
     (c) Recompute the distance matrix after adjustments
  b) If all pairwise distances are greater than or equal to *min_distance*, break the loop and return the updated spatial coordinates of cells
- **Output:**
  - Adjusted spatial coordinates of cells $(x', y')$.

#### 3 Experimental Details

##### 3.1 SpaTedSim simulation procedure and settings

(1) Varying variables in the simulation include:
Migration rate: $\mu_m$ = [0,0.25,0.5,0.75].
Dropout: 0 or 25%.
Number of cells: 1024 or 4096.
(2) Simulation parameters for simulating gene expressions, lineage barcodes and spatial coordinates using SpaTedSim:

Table 1: Simulation settings for gene expression in SpaTedSim

| | |
|---|---|
| Number of genes (ngenes) | 500 |
| Number of Identity Vectors ($N_{IV}$) | 30 |
| State Identity Vector stepsize (*step*) | 0.5 |
| Maximum number of state shifts for one division (*max_walk*) | 6 |
| Asymmetric division rate (p_a) | 0.6 |
| Identity Vector center (starting value for diff-IF) | 1 |
| Number of diff-Identity Vectors ($N_{diff}$) | 20 |
| nondiff-SIV standard deviation ($\sigma$) | 0.5 |
| Probability of nonzero gene effect (ge_prob) | 0.3 |
| Probability of outlier gene (prob_hge) | 0.03 |
| Mean of capture efficiency $\alpha$ (alpha_mean) | 0.1 |

| | |
|---|---|
| Standard deviation of capture efficiency $\alpha$ (alpha_sd) | 0.02 |

Table 2: Simulation settings for lineage barcodes in SpaTedSim

| | |
|---|---|
| Number of characters | 32 |
| Excision dropout rate | 0 |
| Distribution for mutation sampling | Exponential |

Table 3: Simulation settings for spatial coordinates in SpaTedSim

| | |
|---|---|
| Standard deviation of 2-D Brownian motion($\sigma_2$) | 0.6 |
| Division radius ($r_d$) | 3(1024 cells), 6(4096 cells) |
| Migration radius ($r_m$) | 6(1024 cells), 12(4096 cells) |
| Spatial coordinate range (both x and y) | (-3,3) for 1024 cells, (-6,6) for 4096 cells |
| min_distance | 0.2 |
| max_distance | 1 |
| edge length of the hexagon spots | 0.4 |
| batch_effect_size | 1 (small batch effect), 2 (large batch effect) |

##### 3.2 Detailed settings of running TemSOMap on SpaTedSim datasets

We use the same default settings to test the performances of TemSOMap on the simulated datasets (1024 cells and 4096 cells, with and without dropouts in the lineage barcode data). Here we give the default parameter settings for TemSOMap. In Table 4, the first group shows the parameters for preprocessing lineage barcodes to obtain lineage tree and clones, and the second group shows the parameters for the mapping algorithm in TemSOMap.

The first three parameters (weights for calculating Hamming distance) are default values used in the Neighbor-Joining method, DCLEAR. To obtain cell clonal information, we set the number of clones to be 4 (discussed in the main manuscript). Regarding hyperparameters used in TemSOMap loss function, the expression similarity loss terms have the highest weight ($\lambda_1$), the terms involving cell lineage information have relatively high weights but lower than the expression similarity terms ($\lambda_2$). Weight for location-aware entropy loss ($\lambda_3$) is relatively high because the magnitude of values of this loss term is very low.

For SpaTedSim-fitted STARmap mouse cortex data, all parameters are kept the same as the other simulated datasets, except for the weight for variance loss due to the different magnitude of the spatial coordinates ($\lambda_4$, ranging from 0.001 to 0.0000001). The visualizations and benchmarks are reproducible using the notebooks and source codes at: `https://github.com/ZhangLabGT/TemSOMap`.

##### 3.3 Detailed settings of running TemSOMap on E9.5 mouse embryo data

For this experiment, we perform preprocessing of the gene expressions using Scanpy. First, we perform filtering on the input expression data to remove the mitochondrial genes and lowly expressed cells/genes (min_genes = 10 and min_cells = 3). Second, we perform the log transform on the counts. Then, we identify the top 600 highly variable genes from the data. From these genes, 593 genes are present in both Stereo-seq data and the lineage-traced scRNA-seq data. We use 400 of the 593 genes as the training gene set for

Table 4: Parameter settings for TemSOMap on SpaTedSim datasets

| | |
|---|---|
| Weight for unmutated state for Hamming distance | 1 |
| Weight for mutated state for Hamming distance | 4 |
| Weight for dropout state for Hamming distance | 1 |
| Number of clones | 4 |
| Weight for spot-wise expression loss ($\lambda_1$) | 1 |
| Weight for gene-wise expression loss ($\lambda_1$) | 1 |
| Weight for lineage loss ($\lambda_2$) | 0.01 |
| Weight for clonal loss ($\lambda_2$) | 0.01 |
| Weight for location-aware entropy loss ($\lambda_3$) | 0.1 |
| Weight for variance loss ($\lambda_4$) | 0.001 |
| Learning rate | 0.001 |
| Number of iterations | 4000 |

TemSOMap as well as other methods, and use the other 193 genes to test TemSOMap's ability to predict the spatial distribution of unseen genes. For hyperparameters in TemSOMap, all parameters are kept the same as the previous tests (Table 4), except for the weight for variance loss due to the different magnitude of the spatial coordinates, which we set to $\lambda_4 = 0.00001$. The visualizations are reproducible using the notebooks and source codes at: `https://github.com/ZhangLabGT/TemSOMap`.

##### 3.4 Running state-of-the-art methods and evaluation metrics

In order to benchmark the performance of TemSOMap, three other existing methods that are able to infer spatial coordinates of cells from gene expression and spatial transcriptomics data were implemented: Tangram, CeLEry, and SpaOTsc.

For Tangram, we refer to the github repo (`https://github.com/broadinstitute/Tangram`) tutorial and use the default parameters to run. For optional density prior, we provide the true density information from SpaTedSim. For a fair comparison, we are not using the annotation of cell clusters as an input for Tangram. For the mouse embryo data, in order to obtain single-cell level spatial coordinates, we adopt the similar manner with TemSOMap, which takes the product of the mapping matrix and the spot coordinates $MS$ to get the single-cell coordinates.

For both CeLEry (`https://github.com/QihuangZhang/CeLEry/blob/main/tutorial/tutorial.ipynb`) and SpaOTsc (`https://github.com/zcang/SpaOTsc/blob/master/short_tutorial/spaotsc_tutorial_short.ipynb`), the tutorial provided by the authors of each method was downloaded and used as a guide. Our datasets were normalized and packaged into the AnnData format CeLEry takes as input; they were then used to train the CeLEry model and evaluate its prediction of the coordinates of our test dataset. SpaOTsc required additional preprocessing on the datasets, which was done using Scanpy. After normalization, the SpaOTsc cost matrix was built using the Matthews correlation coefficient between binarized expression and spatial count data. This cost matrix, along with both binarized and non-binarized data, was used to generate the SpaOTsc transport plan, which served as a mapping between individual cells and spots.

Four different metrics were used to compare TemSOMap to these existing methods. Two of them are the average MSE and MAE between predicted and ground truth coordinates for each cell. For the other metrics, we first computed pairwise distance matrices between individual cells; one for predicted coordinates and one for the ground truth. The third metric used is the Pearson's correlation coefficient between the two pairwise distance matrices. Finally, we found the 20 nearest neighbors for each cell based on the pairwise distance matrices. We then found the Jaccard index between the predicted and ground truth nearest neighbors of each cell; the average of these values served as our fourth metric.

#### References


1. Chan, M.M., Smith, Z.D., Grosswendt, S., Kretzmer, H., et al.: Molecular recording of mammalian embryogenesis. Nature **570**(7759), 77–82 (Jun 2019). https://doi.org/10.1038/s41586-019-1184-5, https://doi.org/10.1038/s41586-019-1184-5
2. Dries, R., Zhu, Q., Dong, R., Eng, C.H.L., et al.: Giotto: a toolbox for integrative analysis and visualization of spatial expression data. Genome Biology **22**(1) (Mar 2021). https://doi.org/10.1186/s13059-021-02286-2, http://dx.doi.org/10.1186/s13059-021-02286-2
3. Pan, X., Li, H., Zhang, X.: Tedsim: temporal dynamics simulation of single-cell rna sequencing data and cell division history. Nucleic Acids Research **50**(8), 4272–4288 (Apr 2022). https://doi.org/10.1093/nar/gkac235, http://dx.doi.org/10.1093/nar/gkac235
